# Nascent RNA sequencing identifies a widespread sigma70-dependent pausing regulated by Gre factors in bacteria

**DOI:** 10.1101/2020.10.25.354225

**Authors:** Zhe Sun, Alexander Yakhnin, Peter C. FitzGerald, Carl E. Mclntosh, Mikhail Kashlev

**Affiliations:** RNA Biology Laboratory, National Cancer Institute, National Institutes of Health, Frederick, MD 21702, USA; Genome Analysis Unit, National Cancer Institute, National Institutes of Health, Bethesda, MD 20892, USA

## Abstract

Promoter-proximal pausing regulates expression of many eukaryotic genes and serves as checkpoints for assembly of elongation/splicing machinery. Little is known how broadly the pausing is employed in transcriptional regulation in bacteria. We applied NET-seq combined with RNase I footprinting for genome-wide analysis of σ^70^-dependent transcription pauses in *Escherichia coli*. Many *E. coli* genes appear to contain clusters of strong backtracked pauses at 10-20-bp distance from the transcription start site caused by retention of σ^70^ subunit in RNA polymerase. The pauses in 10-15-bp register of the promoter are dictated by binding of σ^70^ to canonical −10 element, 6-7 nt spacer and “YR_+1_Y” motif centered at transcription start site all characteristic for strong *E. coli* promoters. The promoters for the pauses in 16-20-bp register contain an additional −10-like sequence positioned on the same face of the DNA duplex as the original −10 element suggesting that σ^70^ hopping was responsible for these pauses. Our *in vitro* analysis reveals that RNA polymerase backtracking and DNA scrunching are involved in these pauses that are relieved by Gre transcript cleavage factors. The genes coding for transcription factors are enriched in these pauses suggesting that σ^70^ and Gre proteins regulate transcription in response to changing environmental cues.

## INTRODUCTION

Transcription pausing is a fundamental mechanism shared by all kingdoms of life and known to regulate gene expression, alternative splicing, co-transcriptional RNA processing, termination, and synchronize transcription and translation^1–4^. In *E. coli*, a special pausing signal G_−10_Y_−1_G_+1_ (Y_−1_ represents the pause site) was identified^5–7^, which slows down RNA polymerase (RNAP) near a translation start site allowing coordination of RNAP movement with co-transcriptional translation. The elemental pause could be further stabilized by an RNA hairpin formed in the RNA exit channel of RNAP^8,9^ or by RNAP backtracking^10,11^. During backtracking, RNAP moves backward along the DNA and the nascent RNA leading to extrusion of the RNA 3’ end into the RNAP secondary channel to induce long pause or transcription arrest^10,12^. The backtracked pauses can be rescued by removing the extruded 3’ RNA end in a cleavage reaction stimulated by Gre cleavage factors^13–15^. In addition, transcription factors, such as RfaH and RpoD (σ^70^) have been shown to induce transcription pausing by interacting with RNAP and DNA^4^.

The housekeeping initiation factor sigma70 (σ^70^) recognizes the −10 and −35 elements in the promoter regions to form an open promoter complex (RPo) by unwinding DNA duplex between the −10 element and the transcription start site (TSS)^16^. Normally, escape of RNAP from the promoter causes release of σ^70^ in a stochastic manner^17^. However, *in vivo* ChIP-seq and *in vitro* biochemical data showed that σ^70^ could be retained in RNAP at a significant distance from the promoter and the efficiency of retention depended on the transcription unit^18–20^. The −10-like sequence in the initial transcribed region of the λpR’ promoter has been shown to cause retention of σ^70^ in Eσ^70^ holoenzyme leading to transcription pausing^21^. In the σ^70^-dependent pause, the DNA strands in the transcription bubble become scrunched inside RNAP and the strain accumulated during scrunching results in a backtracked σ^70^-dependent pause state^22–24^ allowing proper loading of the accessory antitermination bacteriophage λ Q protein. Elongation factors GreA and GreB release σ^70^-dependent pauses *in vitro*^14,25,26^ by stimulating the nascent RNA cleavage in backtracked RNAP. Although the σ^70^-dependent pauses have been detected at several *E. coli* and phage promoters^27,28^, their robustness, prevalence and their effect on gene expression *in vivo* remain largely unknown.

Nascent elongating transcript sequencing (NET-seq) has been developed to monitor the genome-wide transcription pausing at single nucleotide resolution *in vivo*^29^. In this study, we reported a modified version of NET-seq combined with RNase I footprinting of the nascent transcripts (RNET-seq) for genome-wide identification of σ^70^-dependent transcription pauses in *E. coli*. We found that a strikingly large number of *E. coli* genes contain strong σ^70^-dependent pauses in 5’ UTRs clustered at 10-20-bp distance from promoters. We determined the DNA signals essential for these pauses, identified the mechanism of their rescue by Gre factors, and proposed their role in gene repression and transcription response to changing environmental cues.

## RESULTS

### RNET-seq identifies robust σ^70^-dependent transcription pausing in *E. coli*

In this work, we employed the RNET-seq technique for genome-wide identification of paused ternary elongation complexes (TECs) of RNAP containing σ^70^ subunit isolated from WT and Δ*greAB E. coli* cells (Fig. 1a). Briefly, transcription-engaged RNAP was released from *E. coli* nucleoids by treatment with DNase I and RNase I followed by immobilization on Ni^2+^-NTA agarose beads through His-tag fused to σ^70^ (RpoD) or the β’ (RpoC) subunit (σ^70^ and β’ datasets). Treatment with RNase I degraded all transcripts except for the 3’-proximal fragments strongly protected by RNAP. The immobilized complexes were capable of [α-^32^P] UTP incorporation and susceptible to the RNA cleavage stimulated by GreB indicating their engagement in active transcription (Fig. 1b). A strong positive correlation between the biological replicates of RNET-seq was observed by comparing the normalized counts of reads in each gene (Fig. 1c; Extended Data Fig. 1). The *in vitro* RNase I footprints of the regular paused TECs containing σ^70^ subunit consisted of 16-17 nt of the 3’-proximal RNA (Extended Data Fig. 2). Similarly, the *in vivo* footprints by RNET-seq centered at 17-18 nt and 16-17 nt lengths in β’-WT and σ^70^-WT datasets, respectively (Fig. 1d). We noted that the σ^70^-WT data also contained the short 6-15-nt RNAs deriving from pausing close to promoters. The majority of short 6-11-nt RNAs could not be uniquely mapped to the reference *E. coli* genome and were discarded (Extended Data Fig. 3).

**Fig. 1.**
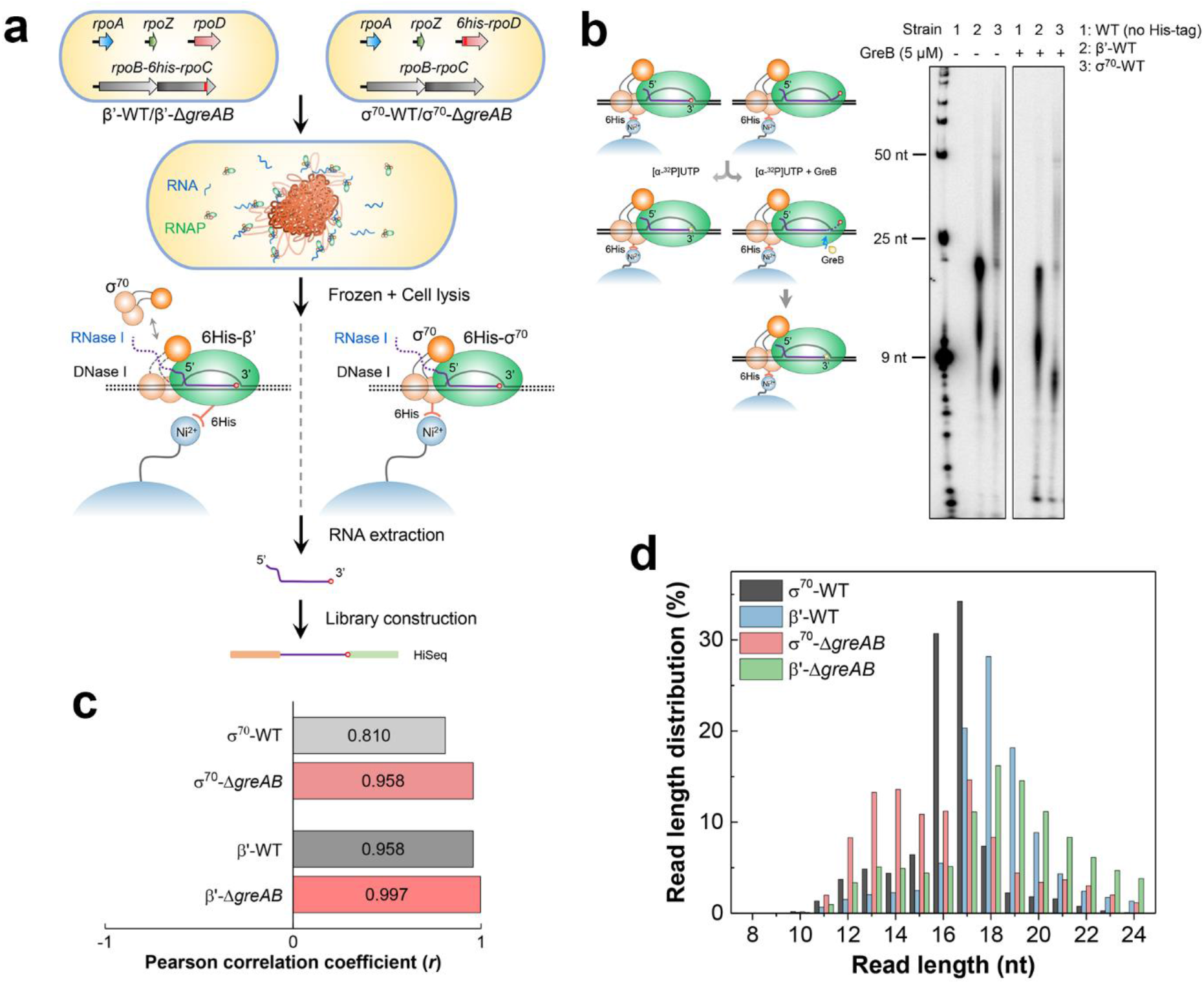
RNET-seq identifies σ^70^-dependent pauses and the corresponding translocation states of RNAP in WT and Δ*greAB* cells. **a,** Principles of RNET-seq. The σ^70^ and β’ strains with 6His-tagged RpoD (σ^70^) and RpoC (β’) subunits were used for purification of the intact paused TECs from bacterial nucleoids after treatment with nucleases (see details in Methods). The green oval represents the RNAP core enzyme. Three connected brown circles represent individual domains of σ^70^ subunit bound to the −10/-35 promoter elements or to RNAP core. **b,** PAGE of the ^32^P-RNA-labeled paused complexes and their sensitivity to the cleavage by GreB protein. WT, *E. coli* W3110 strain lacking His-tag in σ^70^ and β’. The first lane, RNA ladder. **c,** Pearson correlation coefficient (*r*) between two biological replicates from the indicated strains. **d,** Histogram shows RNA length distributions (RNA footprints) for the uniquely mapped RNET-seq reads from the indicated strains. The length of protected RNA allowed determination of translocation register of RNAP in β’-WT at each pause’ 16-17-nt, 18-nt and >18-nt RNAs corresponded to the post-translocated, pre-translocated and backtracked states, respectively (A. Yakhnin *et al*., unpublished observation). The average read lengths for σ^70^-WT, β’-WT, σ^70^-Δ*greAB* and β’-Δ*greAB* strains are 16.3-nt, 18.0-nt, 16.0-nt and 18.3-nt. The <16-nt RNAs derived from pausing at a short <16-bp distance from promoters where the nascent transcripts were not yet accessible to the nucleases.

GreA and GreB proteins were previously identified as the major regulators of RNAP pausing and arrest close to promoters^25,30^. The σ^70^ data from *ΔgreAB* cells showed a characteristic shift of the RNA length from 16-17 nt to 12-17 nt suggesting that, in the absence of Gre factors, σ^70^-dependent pauses predominantly occurred in the 12-17-nt registers downstream from TSS (Fig. 1d, black and pink columns). The σ^70^-dependent pauses in Δ*greAB*, but not in WT cells were also enriched in >17-nt reads arguing that Gre factors efficiently suppressed RNAP backtracking caused by σ^70^, and/or rescued the backtracked complexes^14^ (Fig. 1d, pink column).

### Proximity of σ^70^-dependent transcription pauses to promoters

The peak representing a typical σ^70^-dependent pause in σ^70^-Δ*greAB* cells is shown in Fig. 2a. In total, we identified 7412 and 3543 pauses, whose read counts are at least 20-fold over the median value of all RNA reads in 51-bp window centered at the peak, in σ^70^-WT and β’-WT cells, respectively (Supplementary Table S1). The σ^70^-WT library had lower background than the β’-WT of RNA reads in this 51-bp window, which resulted in a larger number of the pauses counted in σ^70^-WT compared to β’-WT cells. This pattern strongly indicated that the majority of σ^70^ subunit was bound to RNAP within the narrow promoter-proximal regions of the genome. About 26% of the β’-WT pause sites were shared with σ^70^-WT cells and fraction of the shared pauses increased to 57% in Δ*greAB* cells (Fig. 2b). The total number of pauses is also 1.7-1.8-fold higher in Δ*greAB* cells: 12211 pauses in σ^70^-Δ*greAB* cells versus 6498 pauses in β’-Δ*greAB* cells (Supplementary Table S1). These data strongly argued that a substantial fraction of σ^70^-dependent pauses was suppressed or released by Gre factors in WT cells. A large fraction of these pauses resided in UTR and antisense regions (σ^70^ pauses in Fig. 2c; β’ pauses in Extended Data Fig. 4).

**Fig. 2.**
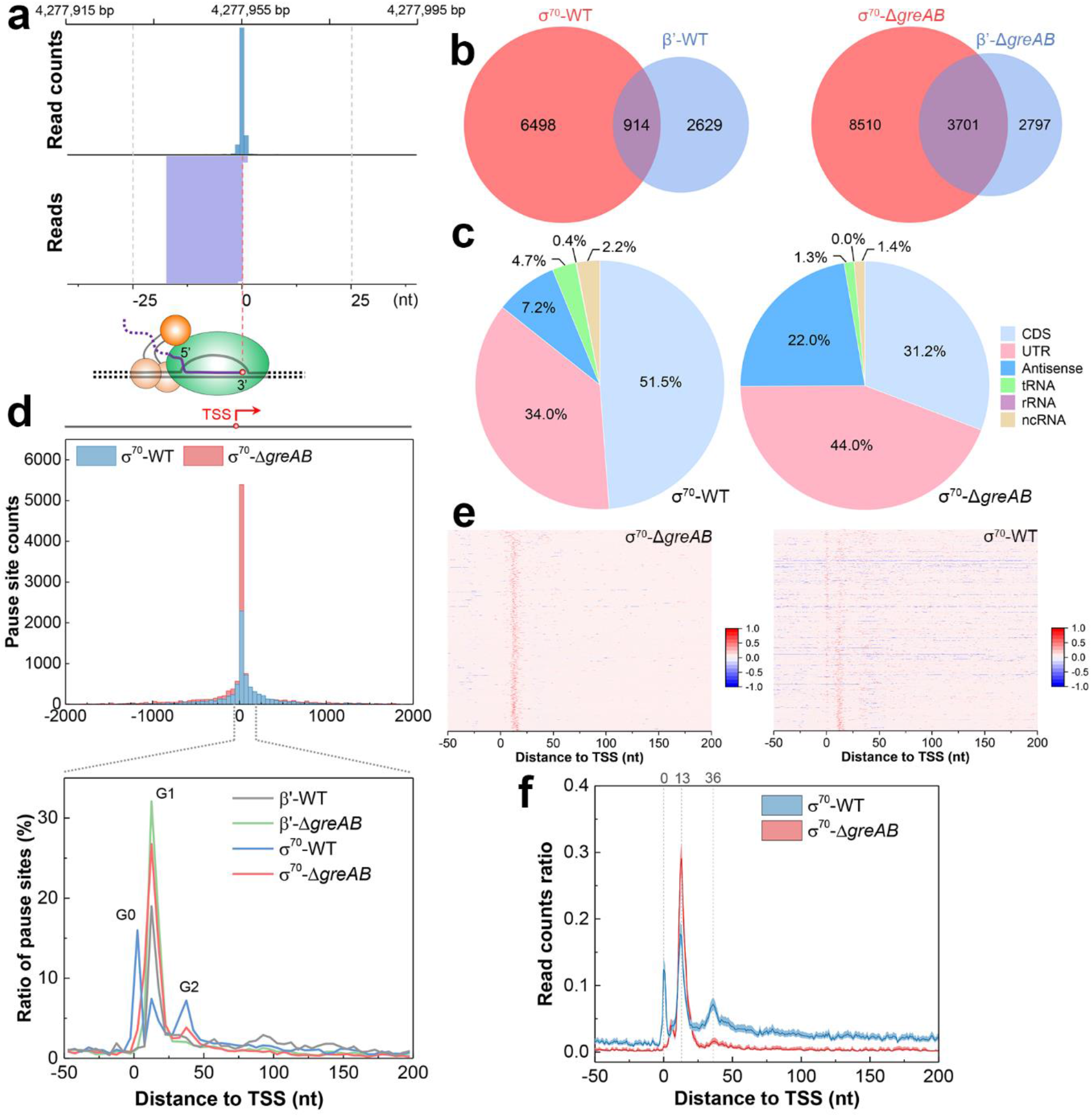
Classification of σ^70^-dependent transcription pauses. **a,** Example of σ^70^-dependent pause upstream of the *yjcE* gene identified by RNET-seq in the σ^70^-Δ*greAB* strain. The genomic coordinates for 3’ ends of all uniquely mapped RNA reads (bottom lane) were determined and the read count for each 3’ end position was calculated and plotted (top lane). The genomic positions where 3’ end/3’ end median (51-bp window) read counts ratio (pause score) was ≥ 20 and read counts/10^6^ reads was ≥ 10 satisfied our stringent definition for a pause site. **b,** Venn diagrams show the total and shared numbers of pauses identified in σ^70^-WT (n = 7412), β’-WT (n = 3543), σ^70^-Δ*greAB* (n = 12211) and β’-Δ*greAB* (n = 6498) strains. **c,** Distribution of σ^70^-dependent pauses among CDS, UTR, Antisense, tRNA, rRNA and ncRNA transcription in σ^70^-WT and σ^70^-Δ*greAB* strains. The “Antisense” pauses included those in CDS, tRNA, rRNA and ncRNA genes. **d,** Distribution of pause sites in promoter-proximal regions. The TSS coordinates identified by dRNA-seq^58^ were used to plot pause counts against the pause distance from the nearest TSS on the same DNA strand. The zero and positive coordinates correspond to the pauses overlapping the TSS or located downstream of the TSS, respectively. The upper panel shows the counts of pauses in 50-nt bins within −2000/+2000-bp window centered at TSS. The bottom panel shows the ratio obtained by dividing count of pause sites in 5-bp sliding window to the total count of pause sites in −50/+200-bp register surrounding TSS. Heatmap **(e)** and mean **(f)** of the read counts for σ^70^-Δ*greAB* G1 pause sites (n = 3099) in *σ^70^-ΔgreAB* (left) and σ^70^-WT (right) strains. The pause sites were ranked based on the pause score (described in **a**). The counts of read aligned to the sense and antisense strands in each coordinate were normalized to 0 to 1 and 0 to −1 by dividing the maximum read count in each −50/+200-bp region. The regions with multiple pause sites were counted only once. **(e)**. The dash line and number on the top indicate distance of the peak from TSS. The line and the shadowed region represent the mean and 95% confidence interval for the read counts ratio **(f)**.

The majority of strong σ^70^-dependent pauses in WT and Δ*greAB* cells was localized within ~50 bp distance downstream of the annotated TSS (Fig. 2d, top). We arbitrarily separated these pauses into G0, G1 and G2 groups located at −2 to 3, 10 to 20 and 31 to 39 bp distance from the closest TSS, respectively (Fig. 2d, bottom). Although these three groups were similarly populated in σ^70^-WT cells, the G1 pauses (Supplementary Table S2) dominated in Δ*greAB* cells arguing that Gre factors primarily suppressed or released pausing at a short 10-20 nt distance from TSS. Heatmap analysis further revealed that G1 pauses were significantly stronger in σ^70^-Δ*greAB* compared to σ^70^-WT cells (Fig. 2e). In contrast, G0 and G2 pauses predominantly observed in σ^70^-WT cells were merely as strong as G1 pauses (Fig. 2f). Notably, G0 pauses had their 5’ RNA ends residing upstream from the closest TSS indicating that they originated from upstream promoters. Most G0 and G2 pauses were substantially weaker than the G1 pauses in β’ cells, and these pauses were not analyzed any further.

### Two categories of G1 pauses

As reported previously, a promoter-like −10 sequence located downstream from the original promoter was essential for σ^70^-dependent pausing^21,27,28^. To investigate whether the −10-like region (−10LR) was involved in G1 pauses in σ^70^-Δ*greAB* cells, we sorted these pauses based on their distance from the TSS and aligned them centering at the corresponding TSS. The putative −10LR was identified for the pauses in 16-20-nt, but not in the 10-15-nt G1 register (Extended Data Fig. S5). Information content (Ri) quantification of −10LR by a σ^70^ model^31^ showed an average Ri above 0 for pauses in 16-20-nt from the TSS (Fig. 3a). Based on this difference, G1 pauses were divided into two categories: proximal G1p (10-15 nt) and distal G1d (16-20 nt), the latter showed significantly higher Ri of −10LR compared to all σ^70^ promoters from RegulonDB^32^ (Fig. 3b). The significantly shorter read length at G1p pauses compared to G1d and all other peaks indicated a close proximity of G1p pauses to promoters with their 5’ end residing directly at the TSS (Fig. 3c). Accordingly, the relatively long read length of G1d pauses suggested high fraction of backtracked pausing (Fig. 3d).

**Fig. 3.**
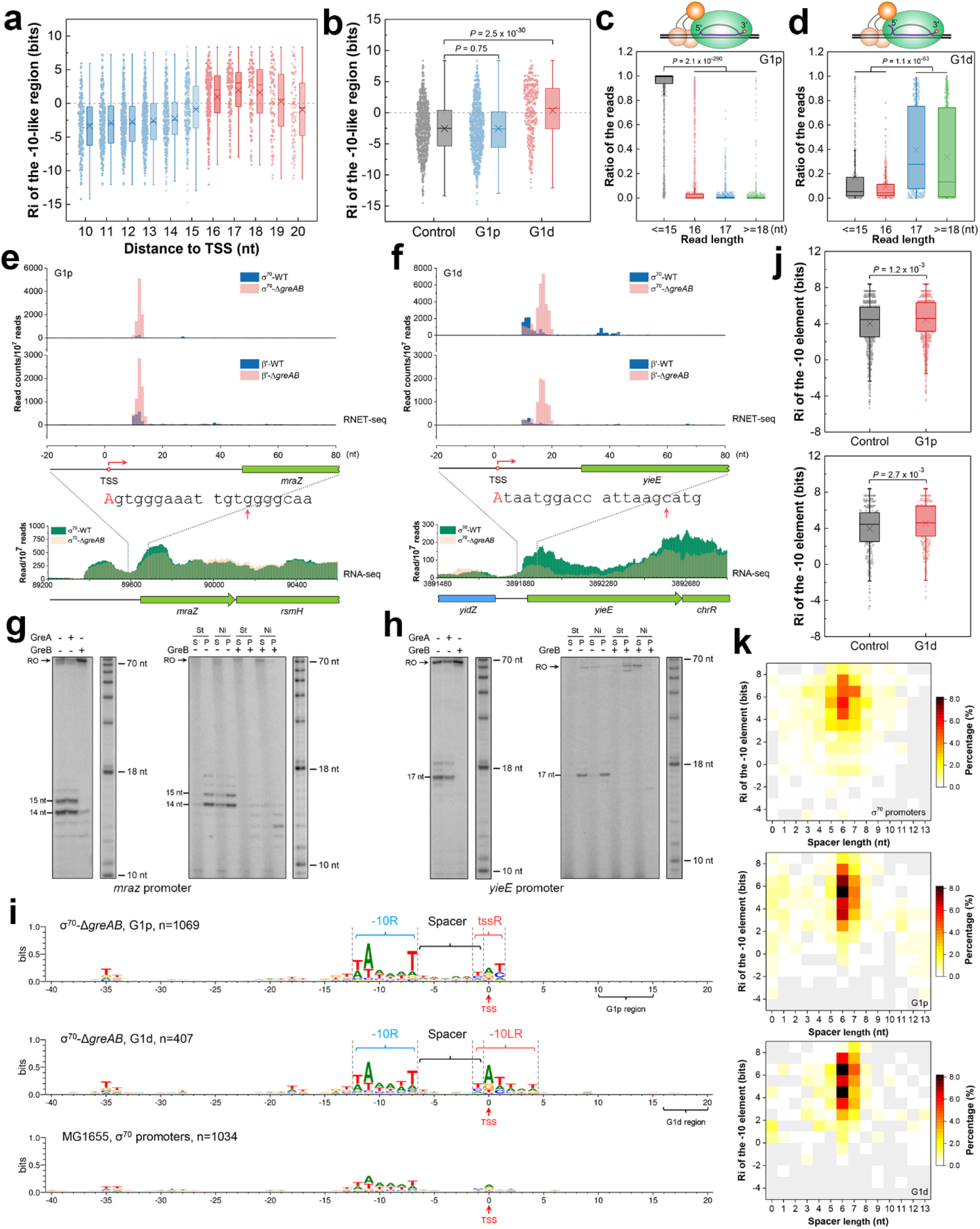
Statistical and *in vitro* biochemical analysis of G1 pauses. **a,** Information content (Ri) for −10LR (−10-like region) encoded by all σ^70^-Δ*greAB* G1 pauses as a function of its distance from TSS. The second base in the −10-like hexamer marked the location of the −10LR. The highest Ri of the hexamers ranging from −1 to +2 was adopted and assigned to −10LR (n = 3099). **b,** Boxplot compares the Ri of −10LR for proximal G1p and distal G1d pauses. All σ^70^ promoters from RegulonDB with a labeled −10 element were used as a control (n = 950). **c, d,** Read length distribution at G1p and G1d pauses, respectively. Ratio of reads, number of reads with specific length(es)/number of total reads. The cartoon on the top depict the backtracked translocation states of G1d complexes based on a significant difference of their read lengths. Note, that the short ≤15-nt RNAs detected at most G1p pauses were due to the close proximity of G1p pauses to TSS that precluded determination of translocation state of G1p complexes by treatment with RNase I. **e, f,** RNET-seq and RNA-seq profiles of two representative genomic regions containing G1p and G1d pauses identified by RNET-seq at *mraZ* and *yieE* promoters. The first 20 nt of *mraZ* and *yieE* transcripts are shown. The red capital letters and arrows indicate the TSS and the pause peaks from RNET-seq data. **g, h,** *In vitro* validation of the σ^70^-dependent G1p/G1d pauses at *mraZ* and *yieE* promoters. The left panel shows nascent RNA in the paused complexes obtained in the presence and absence of GreA or GreB. Immobilization on streptavidin beads through 5’-biotin DNA was used to confirm integrity of the RNA-labeled paused complexes (right panel). Eσ^70^ with His-tagged σ^70^ was used for the assay confirming presence of σ^70^ in the paused complexes. RO, run-off transcripts; St, streptavidin; Ni, Ni^2+^-NTA agarose; S, supernatant; P, pellet. **i,** Sequence logo for *σ^70^-ΔgreAB* G1p and G1d promoters and for σ^70^ promoters from RegulonDB. The DNA sequences were aligned relative to the TSS. Only the strongest pause was used for analysis of the TSSs following multiple pause sites. Coordinate “0” represents TSS (commonly marked as the +1 site) in the sequence logo, otherwise the standard “+1” TSS nomenclature was used. −10R, −10 promoter element; tssR, region surrounding TSS; −10LR, −10-like region; spacer, spacing region between − 10R and TSS. **j,** Boxplot comparing Ri of the −10 elements for G1p (top, n = 1069) and G1d (bottom, n = 407) promoters; −10R of the same numbers of randomly chosen promoters were used as a control. **k,** Heatmap showing correlation between distribution of spacer length and information content (Ri) of the promoter −10 element for all σ^70^ promoters (top), promoters containing G1p (middle) and G1d (bottom) pauses. The two-tailed Mann-Whitney *U*-test was used for the statistical analysis shown above.

Fig. 3e and f show representative G1p and G1d pause sites identified by RNET-seq at *mraZ* (G1p) and *yieE* (G1d) promoters. An *in vitro* transcription assay confirmed the presence of pauses at the same distance from the TSS as the pauses that were determined by our RNET-seq. These pauses were not observed in the presence of GreA and GreB proteins indicating that G1p and G1d pauses included backtracking intermediates rescued by Gre factors (Fig. 3g, h, left). Pulling down the ^32^P-RNA-labeled paused complexes by His-tagged σ^70^ or by biotin group in template DNA confirmed presence of a major fraction of σ^70^ subunit in both paused complexes *in vitro* (Fig. 3g, h, right). The close similarity of the *in vitro* and *in vivo* results strongly argued that σ^70^ subunit was involved in G1p and G1d pauses in promoter-proximal regions of many *E. coli* genes *in vivo*.

An alignment of the G1p and G1d promoter sequences revealed several DNA motifs located immediately upstream from the G1 pauses, which were absent in the reference group of σ^70^-dependent promoters. The distinct promoter −10 element (−10R) for both the G1p and G1d promoters (Fig. 3i) indicated a more conserved −10R and/or more conserved 6-bp spacer length between −10R and TSS for the G1 promoters. A significantly higher Ri of the −10R element was observed at promoters located upstream G1p and G1d pauses (Fig. 3j), and for the entire subset of σ^70^ promoters followed by the pause sites identified in this work (Extended Data Fig. S6). A heatmap of Fig. 3k (top) showed a relatively broad spacer length distribution among all σ^70^-dependent promoters in *E. coli*. In contrast, the G1p (63%) and G1d (66%) promoters had more uniform 6-7 nt spacer between −10R and TSS, indicating that the narrow spacer length might contribute to the strength of G1 pauses (Fig. 3k, middle and bottom). The TSS region (tssR) of G1p promoters consisted of three nucleotides centered at +1 TSS was enriched with a “YR_+1_Y” motif with a +1 purine (R) surrounded by two pyrimidines (Y) (Fig. 3i). Interestingly, the same “YR_+1_Y” motif preceded by 6-nt spacer was previously reported as a strong predictor for genome-wide TSS position and promoter strength^33^. The similar tssR motif was also identified in the reported σ^70^ promoters followed by σ^70^-dependent pauses (Extended Data Fig. 7). Thus, promoter-proximal σ^70^-dependent pausing appears to exhibit two distinct mechanisms involving binding of a σ^70^ to strong canonical −10R promoter element, optimal 6/7-bp spacer, and “YR_+1_Y” tssR (G1p promoter), and those containing an additional −10LR sequence at a conserved 11-bp distance downstream from the −10R of the original promoter (G1d promoter). We noticed that the distance between −10R and −10LR sequences approximately corresponds a single helical turn of B-DNA placing these elements on the same side of the DNA helix, which may facilitate a transition from G1p to G1d pause by σ^70^ hopping (see Discussion for more details).

### Mutational *in vitro* analysis confirms importance of −10R, −10LR, tssR elements and spacer for G1p and G1d pauses

Our *in vitro* testing of several G1p and G1d promoters showed that the pausing patterns and sensitivity to Gre factors closely matched the *in vivo* results (Figs. 4a, b; Extended Data Fig. 8, 9). Point mutations, introduced to increase the Ri of the −10R of these promoters (−10R Ri+), significantly increased G1p, but not G1d pause strength, indicating that strong binding of σ^70^ to - 10R was essential for the G1p pauses (Fig. 4c, d; Extended Data Fig. 10). On the other hand, mutations (−10LR Ri+/−) increasing or decreasing Ri of the distal −10LR of G1d promoters increased or decreased the G1d pause strength, respectively arguing that the downstream −10LR was involved in G1d pausing (Fig. 4e; Extended Data Fig. 11c, d). Y-to-R mutations at “Y_−1_R_+1_Y_+2_” tssR of G1p promoters that decreased their Ri significantly reduced G1p pausing while the R_−1_-to-Y/R_+2_-to-Y mutations increasing Ri of a subset of G1p promoters carrying the sub-optimal R_−1_R_+1_Y_+2_/Y_−1_R_+1_R_+2_ sequence moderately increased the pauses indicating contribution to pause strength of the pyrimidine residues adjacent to TSS (Fig. 4f; Extended Data Fig. 11a, b). Although not all gain-of-function G1d promoter −10R and G1p promoter tssR mutations improved the pause strength (Fig. 4d, f), the statistical analysis of loss-of-function mutations strongly indicated that - 10R and “Y_−1_R_+1_Y_+2_” tssR of G1p promoters, as well as −10R and −10LR of G1d promoters were both essential for G1 pauses.

**Fig. 4.**
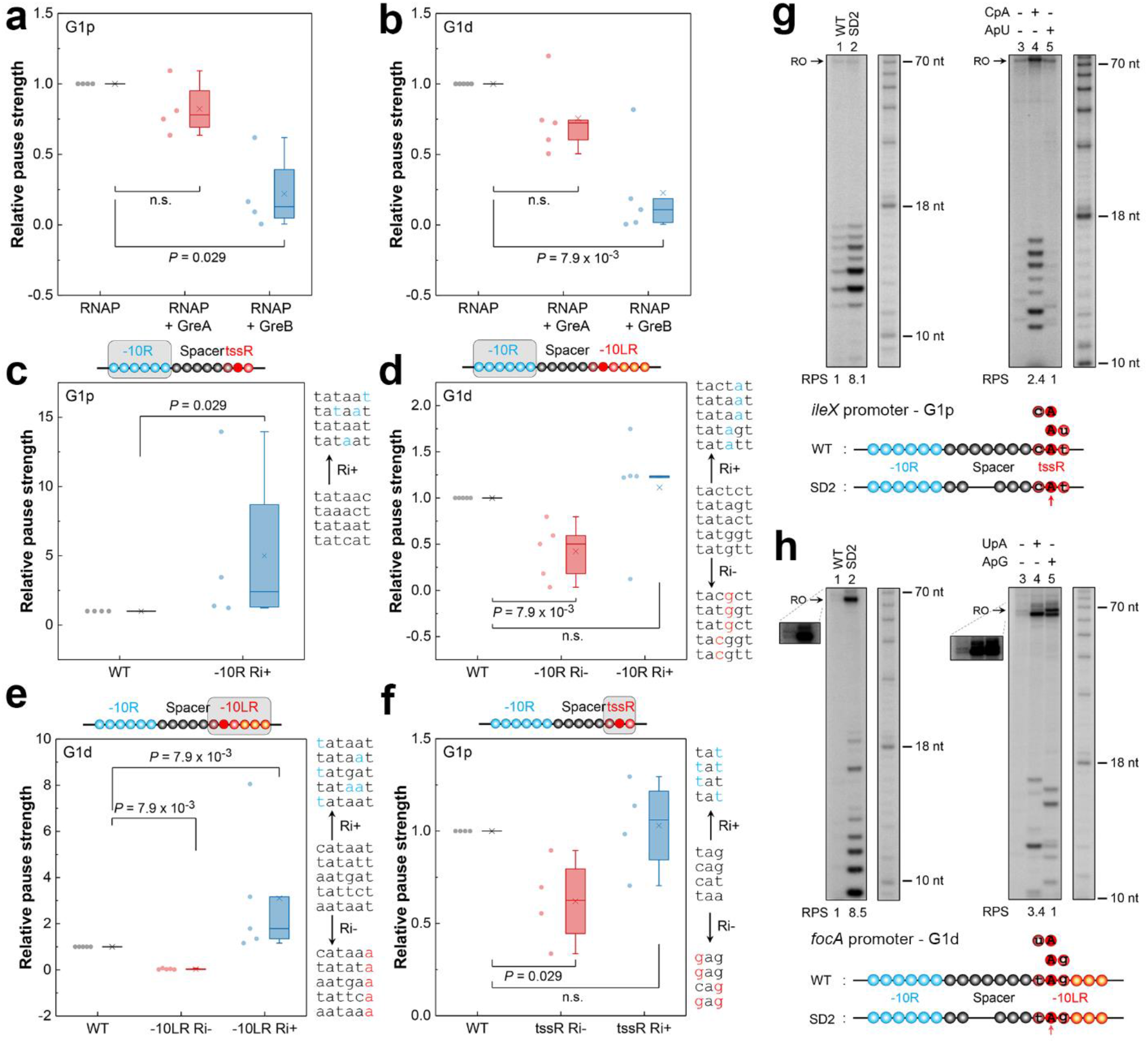
-10R, spacer length and tssR/-10LR determine G1p and G1d pauses *in vitro*. Boxplots of pause strength for G1p **(a)** and G1d **(b)** pauses in the absence and presence of GreA or GreB. G1p (*exuR*, *mraZ*, *ileX* and *mocA*) and G1d (*yieE*, *minC*, *gadW*, *mrdB* and *artP*) promoters were used for the analysis. The pause strength was determined by dividing the signal intensity of run-off and paused RNA products to the signal intensity of paused RNA product in the gel for each *in vitro* template (Pause strength = Signal intensity[paused RNA]/(Signal intensity[paused RNA] + Signal intensity[run-off])). The pause strength in the absence of Gre factors was taken as 1 **(a, b)**. Boxplots show the effect on pause strength of −10R and tssR mutations in G1p promoters **(c, f)**, and −10R and −10LR mutations in G1d promoters **(d, e)**. Pause strength of the WT promoters was set to 1 **(c, d, e, f)**. −10R (−10LR; tssR) Ri-/Ri+, mutated −10R (−10LR; tssR) with decreased or increased Ri are indicated. The grey rectangle in each cartoon represents the motif used for mutation analysis. The original and mutated (colored in blue or red) DNA sequences designed to increase (Ri+) or decrease (Ri-) Ri are shown on the right in gene order. Two-tailed Mann-Whitney *U*-test was used for statistical analysis of the data. Effect of the spacer length on G1p **(g)** and G1d **(h)** pauses. The *in vitro* transcription was initiated on the WT template or on the mutant template with the shortened DNA spacer (left); different dinucleotide RNA primers overlapping the tssR were employed to alter the position of the TSS (right). The inset shows the run-off transcripts with higher exposure to visualize the faint bands. Structural elements of the WT and mutated promoters are shown on the bottom. Each circle represents a single nucleotide. Open blue circles, −10R; Dark red circle, overlapped nucleotide between spacer and tssR/-10LR; Open black and dark red circles, spacer; Red circles, tssR; Red and orange circles, −10LR; Filled red circle, TSS. Red arrows indicate TSS. WT, wild-type promoter; SD2, spacer with 2-nt deletion; RPS, relative pause strength. The analysis included the data from two or more independent experiments.

Finally, we tested an impact of the 6-bp −10R/TSS spacer length on G1 pauses using the *ileX* (G1p) and *focA* (G1d) promoters, both containing the suboptimal 8-bp spacers not typically found in G1 promoters. A 2-bp deletion reducing the *ileX* spacer to 6-bp length caused 8.1-fold increase of the G1p pause strength (Fig. 4g, lanes 1 and 2). Interestingly, a 2.4-fold increase was also observed for the wild-type *ileX* promoter when the regular dinucleotide A_+1_U_+2_ RNA primer corresponding to the native A_+1_ of *ileX* tssR, was replaced with C_−1_A_+1_ primer to induce a 1-bp upstream shift of the TSS, which also shortened the −10R/TSS distance from 8 to 7-bp length (Fig. 4g, lanes 4 and 5; Extended Data Fig. 12). This crucial result strongly indicated that the 6-bp distance between the 5’ RNA end and −10R, rather than the length of the DNA spacer *per se*, was crucial for G1p pauses. A similar result was obtained with the G1d *focA* promoter (Fig. 4h), pointing to a similar role of spacer in both types of G1 pauses. Shortening of the *yieE* promoter spacer from 7 to 6 bp moderately increased the pause strength (Extended Data Fig. 13) indicating that 6-bp spacer length appeared to be optimal for the G1 pauses. Taken together, our mutational analysis confirmed that consensus −10R, 6-bp spacer, and “YR_+1_Y” tssR, all known characteristic for strong *E. coli* promoters, were prerequisites for G1p pausing. In addition, the more distal G1d pauses required the −10LR located at 11-bp distance downstream from the original −10R. This genome-wide result is consistent with the transcription pausing caused by binding of σ^70^ subunit to promoter-proximal −10-like sequences^21,27,28^ that were previously identified at several *E. coli* promoters *in vitro*.

### G1 pauses involve backtracking of RNAP and an extended transcription bubble

To address the structural foundation of G1 pauses, we probed conformational changes in the RNA-DNA hybrid and transcription bubble in TECs at several G1p and G1d pauses identified *in vivo* and confirmed *in vitro*. The complete resistance to RNases T1 and RNase I of the nascent RNA at G1p^*mraZ*^ and G1d^*yieE*^ pauses (in UTRs of the *mraZ* and *yieE* genes) carrying 14-15-nt and 17-nt transcripts, respectively (Fig. 5a, b) and high sensitivity of these complexes to GreB-induced transcript cleavage (Fig. 3g, h) indicated presence of backtracked pauses at both G1 pause sites. Treatment with GreB generated cleavage products shortened by 4-5-nt confirming 4-5-bp backtracking of RNAP at the G1p and G1d pauses (Fig. 5c; Extended Data Fig. S14). As reported earlier, backtracking at ≥ 3-bp distance increases sensitivity to GreB and makes these pauses more resistant to GreA^13^. In contrast, backtracking at 1-2-bp distance makes these pauses more susceptible to GreA^34^. The substantially lower sensitivity of G1p and G1d pauses to GreA compared to GreB (Fig. 4a, b) confirmed backtracking at more than 2-bp distance at the G1 pauses.

**Fig. 5.**
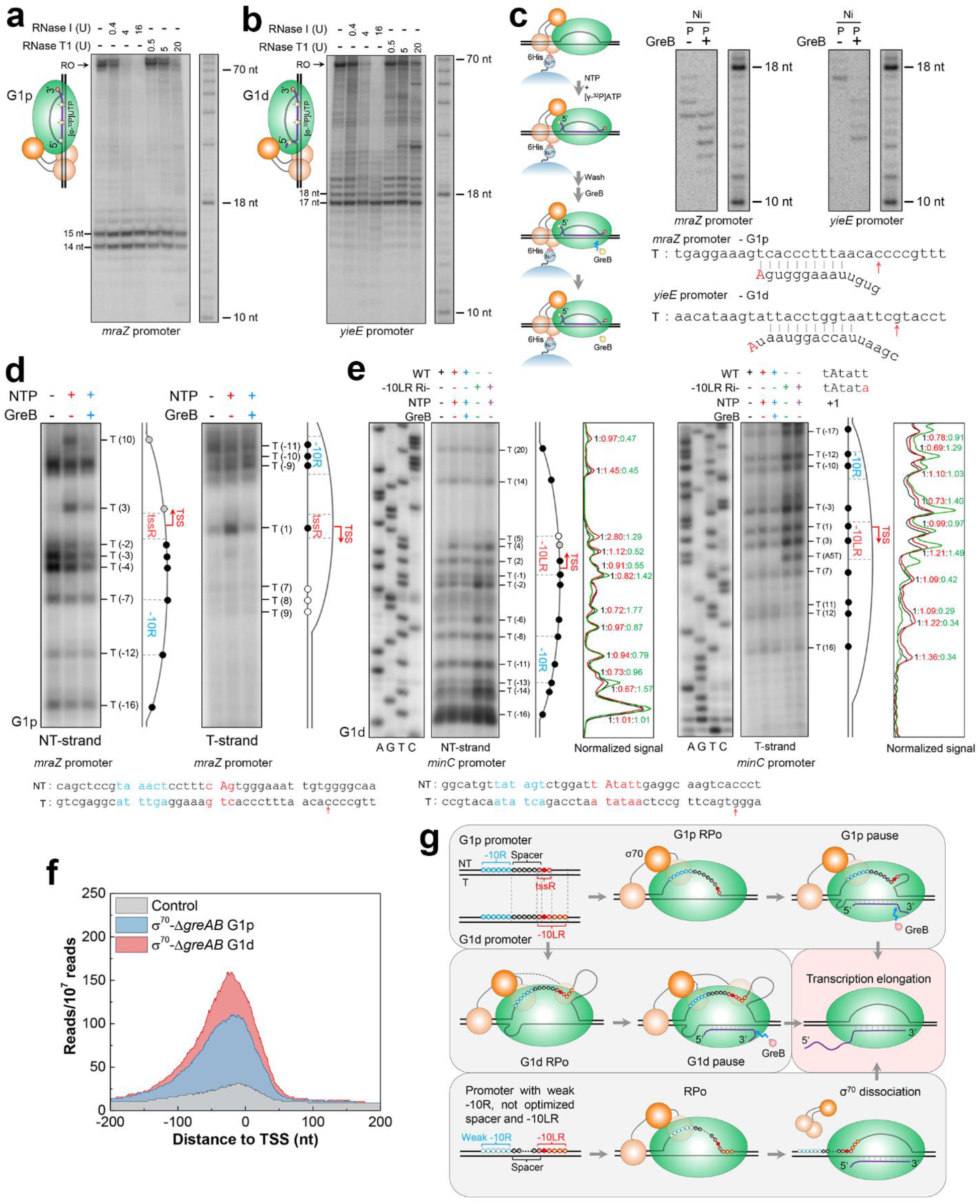
*In vitro* analysis of G1p/G1d pauses and the corresponding open promoter complexes. Protection of the nascent RNA by Eσ^70^ (6His-σ^70^) holoenzyme from digestion by RNases I and T1 at G1p **(a)** and G1d **(b)** pauses. In the regular (non-paused) elongation complex, RNAP protects *in vitro* 14 nt (RNase T1) and 17-18 nt (RNase I) of the 3’ RNA from the nuclease digestion^59^. The cartoons on the left show the proposed alternative translocation states of the RNA in the paused complex. The stars indicate the RNA positions labeled by [α-^32^P] UMP. **c,** GreB-induced transcript cleavage of nascent RNA at G1p (*mraZ*) and G1d (*yieE*) pauses. The workflow for the experiment is shown on the left. Ni, Ni^2+^-NTA beads; P, pellet. The template strand sequences of *mraZ* and *yieE* promoters and backtracked RNAs at the pause sites are shown at the bottom. Red arrows indicate the pausing peaks identified by RNET-seq. **d,** Permanganate footprints of the non-template and template strands of the transcription bubble at the *mraZ* (G1p) promoter. The positions of all T residues in the bubble are indicated. A prominent non-T band between T3 and T10 is a background of permanganate footprinting. The diagrams on the right show the transcription bubble at the *mraZ* promoter during G1p pausing. Black filled circles, T residues sensitive to KMnO_4_ in the absence and presence of NTP; gray filled circles, permanganate-sensitive T residues in the presence of NTP; white filled circle, T residues resistant to permanganate. **e,** Permanganate footprints of transcription bubble at the *minC* (G1d) promoter. Both DNA strands of the *mraZ* and *minC* promoters including the −10R (blue), tssR/-10LR (red) elements and TSS (red capital) are shown at the bottom, and the G1p and G1d pause peaks are marked by red arrows. **f,** Profiles of median ChIP-seq reads coverage at G1p, G1d and control promoters based on the heatmaps (Extended data Fig. 16). **g,** Model depicting the structural properties of σ^70^-dependent G1p and G1d pauses. The interaction of σ^70^ domains with the promoter elements, the DNA scrunching and the corresponding changes in the RNA register at G1p and G1d pauses are indicated.

Potassium permanganate footprinting showed relatively normal size and location of the transcription bubble in the RNAP-promoter open complex (RPo) at the *mraZ* promoter, which codes for a G1p pause (Fig. 5d, −NTP, black lane). In contrast, the TEC at G1p^*mraZ*^ pause showed an unusually long ~26-nt bubble, substantially larger than the bubble detected in the regular TEC carrying the similar length of nascent RNA that was obtained from T7A1 promoter (Fig. 5d, +NTP, red lane; Extended Data Fig. S15). Strikingly, the corresponding RPo and the paused TEC at G1d^*minC*^ promoter exhibited an even larger (>30-nt) transcription bubble not characteristic for the regular RPo and the bubble detected at G1p pause sites (Fig. 5e, −NTP, black lane; and +NTP, red lane). A point mutation introduced to the −10LR of *minC* eliminated the G1d^*minC*^ pause and also reduced the size of the bubble in the RPo^*minC*^ and G1d^*minC*^ pause (Fig. 5e, green and purple lanes). Thus, the extended bubble appeared to be a hallmark of the G1d promoters making them different from the regular and the G1p promoters. Cleavage of the nascent RNA at the G1p^*mraZ*^ and G1d^*minC*^ sites by GreB rescued these pauses (Fig. 3g; Extended Data Fig. 9b). However, treatment with GreB reduced, but did not eliminate the bubble at these pause sites and promoter region (Fig. 5d, e, blue lanes) suggesting that these promoters contained a large fraction of RNAP capable of forming a promoter complex. Presumably, this complex was catalytically inactivated or was elongating the nascent RNA to, but not beyond the G1p and G1d sites in the presence of GreB^30^. Our analysis of the published ChIP-seq/σ^70^ data^35^ confirmed a high enrichment of RNAP holoenzyme in a 400-nt window centered at the TSS of the G1p and G1d promoters compared to the promoters lacking G1 pauses (Fig. 5f; Extended Data Fig. 16).

### σ^70^-induced pausing controls expression of genes involved in transcription regulation

Although regulation of σ^70^-dependent pauses by Gre factors has been well documented *in vitro*^14,27,28^, their biological role and an impact on genome-wide transcription levels warranted further investigation. Our data showed that G1 pauses were significantly increased in number in cells lacking Gre factors. ~70% of all peaks of RNAP enzyme (G1p, 1128/(1128 + 424); G1d, 366/(366 + 158)) identified in Δ*greAB* cells had the matching strong σ^70^ peaks that accumulated at G1 pauses (Fig. 6a), but the RNAP molecules forming these peaks did not proceed into downstream genes. The RNA-seq also showed that genes containing G1 pauses were expressed at a significantly higher level compared to the randomly selected genes (Fig. 6b), which was consistent with the canonical −10 element of the strong G1 promoters (Fig. 3i). Gene ontology (GO) analysis^36^ showed that *E. coli* genes containing the G1 pauses were enriched among genes coding for the general and gene-specific transcription regulators (Fig. 6c). Most important, our RNA-seq analysis of transcription levels in σ^70^-WT and σ^70^-Δ*greAB* cells revealed that genes harboring G1 pauses were consistently downregulated in the σ^70^-Δ*greAB* compared to σ^70^-WT cells, this downregulation was especially pronounced in genes containing the strong G1 pauses (Fig. 6d). Our analysis of the published RNA-seq data revealed that variable environmental stresses counter-regulate transcription of *greA* and *greB* genes (Extended Data Fig. 17). In turn, the G1 pauses are released by either GreA or GreB depending on the type of stress and backtracking distance of the corresponding pause. Thus, our results provided a strong evidence that the highly dynamic G1 pauses with the rapidly exchanging backtracked states are involved in a global regulation of promoter escape and in the local transcriptional networks governed by specialized transcription regulators (Fig. 6e).

**Fig. 6.**
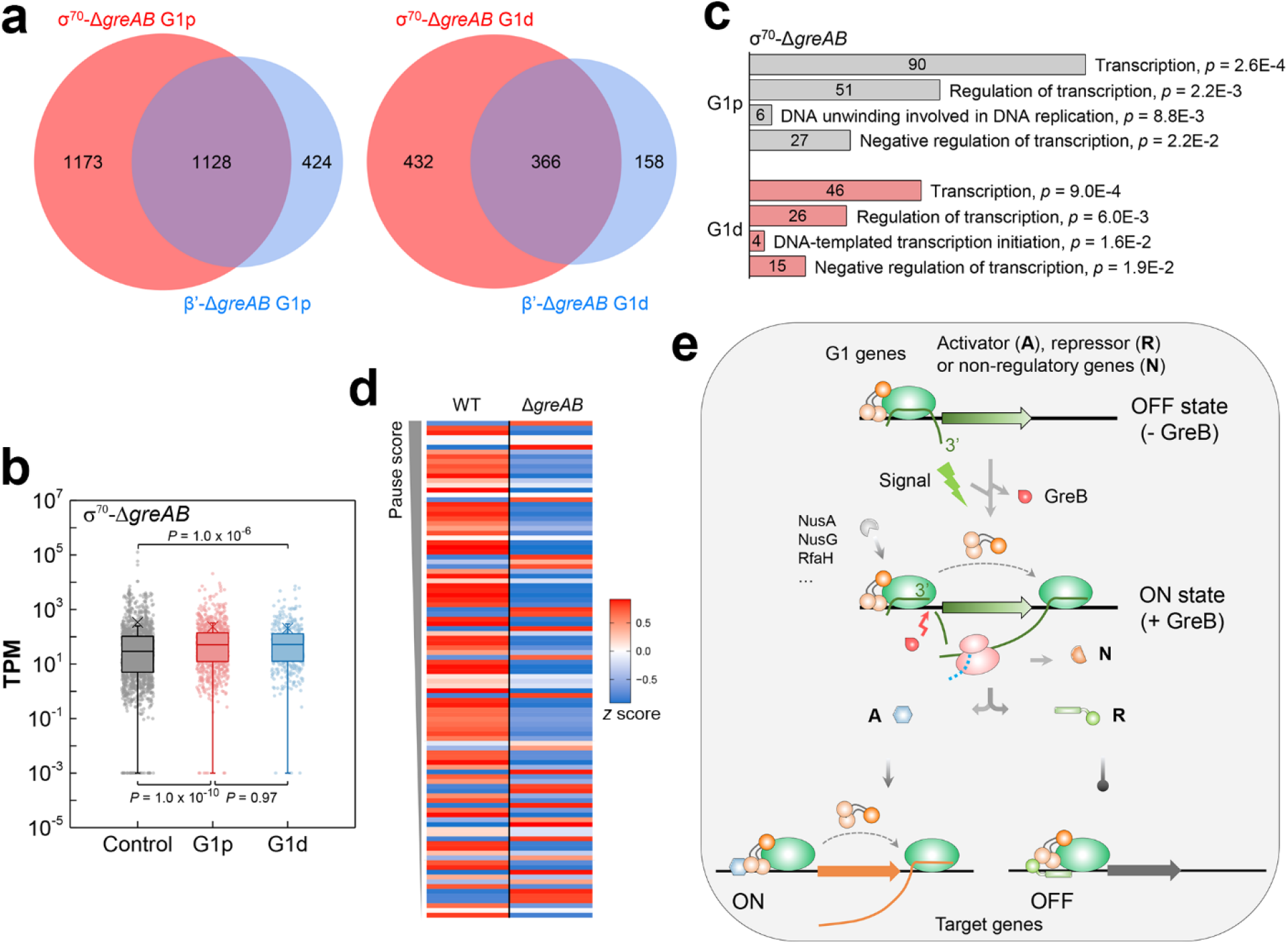
Role of σ^70^-dependent pauses in transcription regulation. **a,** Venn diagrams of all G1p (left) and G1d (right) pauses from Δ*greAB* cells identified in this work. **b,** Boxplot of transcripts per million (TPM) for the genes with and without G1p or G1d pauses. *P* value was calculated by two-tailed Mann-Whitney *U*-test. **c,** Gene Ontology (GO) analysis of genes with G1p and G1d pauses closest to their start codons. All significantly enriched gene categories are listed. The number of genes in each category is shown inside the bars. **d,** Heatmap shows the transcription pattern of genes containing G1 pauses in σ^70^-WT and σ^70^-Δ*greAB* datasets. The G1 genes whose start codon is ≤ 20 bp upstream and ≤ 100 bp downstream from σ^70^-Δ*greAB* G1 pause sites (pause score ≥ 1000, n = 104) are shown. Data from three biological replicates are presented. WT, σ^70^-WT strain; Δ*greAB*, σ^70^-Δ*greAB* strain. **e,** Schematic illustration of the mechanism for σ^70^-induced promoter-proximal pausing, its suppression by GreB, and the impact of the pausing on global regulation of transcription.

## DISCUSSION

Promoter-proximal pausing is broadly employed for regulation of genes in metazoans^1,2,37^. However, only a limited number of bacterial and bacteriophage promoters have been shown to be regulated by promoter-proximal pausing *in vitro*, and the protein factors involved *in vivo* remain unknown. Here, by using σ^70^-based RNET-seq, we identified genome-wide promoter-proximal pauses caused by the σ^70^ subunit in *E. coli* and regulated by Gre factors. We characterized two distinct mechanisms for promoter-proximal pausing that act *in vivo* consecutively at 10-15-bp (G1p) and 16-20-bp (G1d) distances from the TSS. The G1p promoters consist of the canonical - 10R, a 6-7 nt −10R to TSS spacer, and a “YR_+1_Y” tssR. All these features have been previously shown to determine high strength of *E. coli* promoters^33,38–41^. Although the strong σ^70^ binding seemed to facilitate rapid and stable recruitment of RNAP *in vivo*, it also hindered promoter escape due to the strong anchoring of σ^70^ to the canonical promoter elements, ultimately leading to DNA scrunching and RNAP inactivation by backtracking as depicted in Fig. 5g.

The G1d pauses shared a similar promoter-like structure as the G1p pauses but contained an additional −10LR element that causes retention of σ^70^ after RNAP moves further away from promoter. This mechanism is reminiscent of the pauses reported *in vitro* at λpR’ and *lac* promoters^21,27,28^ that also contain −10LR, and to which σ^70^ binds to induce RNAP pausing. We noticed that a large number of G1 promoters contained both of the G1p and G1d pauses, suggesting that the G1p promoters may increase the local concentration of σ^70^ near the promoter DNA to facilitate hopping of σ^70^ to the secondary −10LR sequence located nearby. Indeed, the −10LR of G1d promoters appeared to be positioned at ~11-bp distance downstream from the original −10R on the same face of the DNA helix, which may further promote the σ^70^ hopping. Although elemental pauses have been reported as precursors for the longer hairpin-dependent and backtracked pauses in *E. coli*^4,8^, we could not find the putative elemental pause motifs near the G1 pauses, indicating that they seem not to be essential for σ^70^-dependent pausing.

The metadata analyses (Fig. 5f; Extended Data Fig. 16) showed that the G1 promoters, albeit effectively recruiting RNAP, strongly limit its escape to productive elongation. Holding RNAP at the promoter should block access of other RNAP molecules to the corresponding gene^42,43^, thus, turning RNAP itself to a general transcription repressor. The G1 pausing may represent yet another layer of gene repression in addition to the well-known mechanisms of promoter occlusion by repressors that block open complex formation^38^. In addition, the G1 pauses may expedite transcriptional response to changing environmental cues after being released by Gre factors. Indeed, transcription of the *greB* gene appeared to be induced under different stress conditions (Extended Data Fig. 17) supporting its crucial role in stress response. This mechanism seems to be similar to the robust promoter-proximal pausing of RNA polymerase II and its rescue by TFIIS for rapid response to external signals in eukaryotes^1,2,44^. We found that G1 pauses are enriched in the genes coding for transcription regulators (Fig. 6c) ultimately placing the G1-containing genes to the key nodes involved in regulation of cellular responses to environmental perturbations.

Binding sites for σ^70^ on RNAP core overlap with those for the general Nus factors (NusA, NusG and RfaH) known to synchronize transcription and translation, control pausing during elongation and processivity of RNAP^45–47^ (see Fig. 6e). The G1 pauses may serve as a checkpoint enabling a temporal assembly of these factors at the promoter to guarantee the subsequent proper readout and regulation by the downstream elongation and termination signals. This notion is consistent with a negative correlation between binding pattern of σ^70^ and binding patterns of NusA and NusG observed by ChIP-seq analysis of promoter-proximal regions^48^. The G1 pausing may also stabilize binding of σ^70^ to RNAP and make transcription of the target genes, such as ncRNA and antisense RNA genes, independent of regulation by Nus and Rho factors^18,49^.

Further analysis is required to investigate role of the robust σ^70^-dependent pausing in transcription elongation at a large distance from promoters including transcription terminators^50^. The high evolutionary conservation of σ^70^ suggests that this pausing mechanism is likely shared by other bacteria. RNA polymerase II initiation factors TFIIB and TFIIE^51–53^, possessing homology with bacterial σ factors, are the likely candidates to regulate promoter-proximal pausing in eukaryotes.

## METHODS

### Bacterial strains and growth conditions

*E. coli* strains β’-WT (W3110 *rpoC*-6×His::*kan*) and β’-Δ*greAB* (W3110 *rpoC*-6×His::*kan greA*::*tet greB*::*amp*) were engineered as was previously described^7^. σ^70^-WT (W3110 6×His-*rpoD*) strain was constructed using a CRISPR-Cas9 system. For the His-tagging, a homologous recombination DNA with His-tag DNA sequence (5’-catcaccatcaccatcac-3’) was inserted 3’ of the G residue of the start codon (AT<u>G</u>) of *rpoD* and the ~1.0 kb surrounding DNA was amplified by overlap PCR and cloned into plasmid pTargeT. After electroporation, the tagged strain was identified by PCR and confirmed by Sanger sequencing. The *greA* and *greB* genes were disrupted by P1 transduction from strain β’-Δ*greAB* to obtain the σ^70^-Δ*greAB* (W3110 6×His-*rpoD greA*::*tet greB*::*amp*) strain. The primers used are shown in Supplementary Table S3. All *E. coli* strains were grown in LB medium (tryptone 10 g l^-1^, yeast extract 5 g l^−1^, NaCl 10 g l^−1^) or on LB plate containing 50 μg ml^−1^ kanamycin, 40 μg ml^−1^ spectinomycin, 50 μg ml^−1^ ampicillin or 12.5 μg ml^−1^ tetracycline when appropriate.

### RNET-seq and data analysis

#### Cell collection, lysis and elongation complexes pull-down

An overnight cell culture was diluted in 100 ml LB medium (OD_600_ = 0.02) and cultured at 37 °C to reach a mid-log phase (OD_600_ = 0.5). To stabilize binding of σ^70^ to RNAP core during TEC purification, low ionic strength conditions (described below) were used throughout the purification protocol. Namely, the cell culture was combined with an equal volume of frozen 2 × crush buffer (20 mM Tris-HCl pH 7.8, 10 mM ethylenediaminetetraacetic acid (EDTA), 100 mM NaCl, 1 M Urea, 25 mM NaN_3_, 2 mM β-mercaptoethanol, 10% ethanol, 0.4% NP40, 1 mM PMSF) and the cells were collected by centrifugation (12000 rpm, 15 min, 4 °C), instantly frozen in liquid nitrogen and placed on ice. The cells were resuspended and lysed by 120 kU Ready-Lyse lysozyme (Lucigen), 400 U RNase I (Invitrogen) and 40 U alkaline phosphatase (NEB) at room temperature for 10 min. The chromosomal DNA was pelleted and treated with 300 U RNase I, 6 U Turbo DNase (Invitrogen) and 100 U DNase I (Roche) by vortexing at room temperature for 10 min. After centrifugation (14000 rpm, 3 min, 4 °C), the supernatant containing the solubilized TECs (~700 μl) was incubated with 200 μl of Ni^2+^-NTA beads for 1 hour at 4 °C with continuous shaking (1000 rpm). The beads were washed 4 times with 1 ml of the wash buffer (20 mM Tris-HCl pH 7.8, 1 M betaine, 5% glycerol, 2 mM β-mercaptoethanol, 2.5 mM imidazole) and 3 times by 1 ml pre-elution buffer (20 mM Tris-HCl pH 7.8, 40 mM KCl, 0.3 mM MgCl, 5% glycerol, 2 mM β-mercaptoethanol, 2.5 mM imidazole). The TECs immobilized on the beads were digested once again with 100 U RNase I, 2 U Turbo DNase and 40 U DNase I in 150 μl pre-elution buffer containing 200 μg ml^-1^ bovine serum albumin for 30 min at room temperature with continuous shaking (600 rpm). The beads were washed 4 times with the wash buffer and loaded onto 0.5 ml Ultrafree-MC centrifugal filters (Millipore). The immobilized material was eluted with the wash buffer containing 0.3 M imidazole. The nucleic acids in the eluates were extracted once with 400 μl phenol:chloroform:isoamylalcohol (PCI; 25:24:1) and once with 300 μl chloroform. The top water phase was collected and mixed with 3 volumes (~1200 μl) of isopropanol. After precipitation at −80°C for 30 min and centrifugation, the nucleic acids pellet was washed by 180 μl of 80% ethanol and air-dried. The pellet was dissolved in 12 μl nuclease-free water. The DNA was removed by 2 U Turbo DNase and 10 U DNase I at 37 °C for 15 min. The residual RNA was extracted by PCI, precipitated by isopropanol and solubilized in 10 μl nuclease-free water.

#### Barcode ligation and reverse transcription

The RNA was ligated to 10.7 pmol barcode DNA linker using 200 U T4 RNA ligase 2 (NEB) overnight at 16 °C. The ligation product was extracted by chloroform, precipitated by isopropanol and solubilized in 10 μl nuclease-free water. Reverse transcription was performed using the RNA-DNA chimera and 3 μM phosphorylated reverse transcription primer in 1 × PrimeScript buffer containing 0.5 mM dNTPs, 5 mM DTT, 0.6 U μl^−1^ SuperaseIn RNase inhibitor (Invitrogen) and 10 U μl^−1^ PrimeScript Reverse Transcriptase (Takara) at 48 °C for 30 min. After 2 U RNase H (NEB) treatment for 15 min at 37 °C, the reaction mixture was separated by 10% Urea-TBE PAGE. The cDNA products at 75-100-bp range were excised from the gel and extracted with nuclease-free water for 10 min at 70 °C. The gel chunks were removed by filtering and the cDNA was precipitated by 3 volumes of isopropanol at −80 °C for 30 min and dissolved in 4 μl nuclease-free water.

#### Circularization, library preparation and Illumina sequencing

The resulting cDNA was circularized by 40 U ssDNA ligase (Lucigen) at 60 °C for 4 h. The circularized DNA was subjected to PCR to generate a sequencing library using Illumina index primers and PrimeSTAR Max DNA polymerase (Takara). The PCR product was loaded and electrophoresed by 8% TBE PAGE. The DNA product excised from the gel was extracted overnight by 680 μl DNA soaking buffer (0.3 M NaCl, 10 mM of Tris-HCl pH 8.0, 0.97 mM EDTA) at room temperature. The DNA library was precipitated by isopropanol, washed once by cold 80% ethanol, air dried and dissolved in 8 μl 10 mM Tris-HCl pH 8.0. The concentration of the library was determined by an Agilent 2100 bioanalyzer. Illumina sequencing was performed by the NIH Intramural Sequencing Center. The DNA libraries were quantified by qPCR, pooled and loaded on an Illumina HiSeq 2500 using 2 × 50 bp paired-end sequencing in rapid run mode.

#### Data analysis

After a quality check, the primer sequence was trimmed from the raw R1 reads and duplicates were removed based on the random barcode. The random barcode was removed and the reads were aligned to the *E. coli* genome NC_000913.2 using Bowtie^54^. The 5’ end coordinates of all uniquely aligned R1 reads, which correspond to 3’ end of RNA, were recorded and the total read counts at each coordinate were determined. The coordinate was picked up and defined as a transcription pause site when its read counts was at least 20-fold of the median read counts in a surrounding 51-nt window size and not less than 10 per million reads.

### DNA templates and *in vitro* transcription

The wild-type promoters from the −80 to +60 region relative to the TSS used for *in vitro* transcription, were amplified by PCR using genome as template and cloned into T-Vector pMD19 (Simple, Takara). Primers containing the mutations were used to PCR the whole derived pMD19 plasmid constructed above. The DNA product was self-ligated using T4 DNA ligase (Invitrogen) and transformed to DH5α competent cells. Mutations were confirmed by Sanger sequencing and the plasmid was used to amplify DNA template for *in vitro* transcription. When appropriate, a 5’-biotin-labeled primer was used to amplify DNA template with biotin labeling at the 5’ end of non-template strand. Primers used to amplify DNA templates are listed in Supplementary Table 3. Single round *in vitro* transcription reactions were performed in transcription buffer (40 mM Tris-HCl pH 8.0, 1 mM dithiothreitol, 0.1 mg ml^−1^ BSA, 10 mM MgCl_2_, 50 mM KCl) in two steps. First, 20 nM linear DNA template and 50 nM Eσ^70^ were mixed and incubated at 37 °C for 10 min to form the open complex. When indicated, 200 nM GreA or 50 nM GreB was added in this step. Then 20 μM GTP, UTP, CTP, 2 μM ATP and 5 μCi [γ-^32^P] ATP (PerkinElmer) were used to start the reaction for 10 min. In the second step, the reaction mixture was chased with the addition of 20 μM ATP and 10 μg ml^−1^ rifampicin for 3 min. The reaction was terminated by adding the same volume of 2 × stop buffer (10 M Urea, 250 mM EDTA pH 8.0, 0.05% xylene cyanol and bromphenol blue) and analyzed on 23% (10’1, acrylamide: *bis*acrylamide) polyacrylamide gel with 7 M urea. All procedures of *in vitro* transcription, the following RNase I, RNase T1 cleavage and GreB stimulated cleavage assays were performed at 37 °C unless indicated otherwise.

For testing RNAP activity before RNET-seq, 10 μl Ni^2+^-NTA beads with ECs were washed three times by 200 μl pre-elution buffer. Then 10 mM MgCl_2_ and 10 μCi [α-^32^P] UTP (PerkinElmer) were added to the beads to elongate the nascent RNAs for 10 min. After washing the beads three times by wash buffer, the ECs were eluted by 10 μl wash buffer containing 0.3 M imidazole. For pull-down experiments, 5’-end biotin labeled DNA template and reconstituted Eσ^70^ (6His-σ^70^) were used. The same *in vitro* transcription was done as mentioned above on 8 μl streptavidin and Ni^2+^-NTA beads. After reaction and spinning down the beads, the top solution was collected (“supernatant” fraction) and the bottom beads were immediately washed three times to stop the reaction. The transcription products were released by heating the beads resuspended by the same volume of stop buffer at 95 °C for 5 min (“pellet” fraction). To initiate transcription by dinucleotide, 200 μM CpA, ApU, UpA or ApG (TriLink) were added during open complex formation. Then 20 μM NTPs, 2 μCi [α-^32^P] UTP and 10 μg ml^-1^ rifampicin were added and incubated for 3 min before stopping the reaction.

### RNase I and RNase T1 footprinting of the nascent RNA

In an 8 μl reaction mixture, 20 nM DNA template and 50 nM reconstituted Eσ^70^ (6His-σ^70^ or 6His-β’) was incubated for 10 min on 8 μl Ni^2+^-NTA beads. Then 20 μM GTP, ATP, 2 μM UTP and 3 μCi [α-^32^P] UTP were added to initiate the reaction at the *rrnB* P1 promoter for 10 min. An additional 20 μM CTP was used for the *mraZ* and *yieE* promoters. After chasing the reaction by 20 μM UTP and 10 μg ml^-1^ rifampicin for 3 min, the beads were washed twice and treated by the indicated amount of RNase I (Invitrogen) or RNase T1 (ThermoFisher) for 10 min at 24 °C. The beads were washed two times and extracted with 3 μl PCI to terminate the reaction.

### GreB cleavage assay

The same reaction on Ni^2+^-NTA beads that was used for the RNase footprinting was pre-incubated to form RPo. Transcription was initiated by adding 20 μM GTP, UTP, CTP, 2 μM ATP and 5 μCi [γ-^32^P] ATP for 10 min. The reaction was chased with 20 μM ATP for 3 min. After washing two times, 50 nM GreB was added for 10 min to induce cleavage of the transcripts. The beads were washed twice to stop the reaction and the products were denatured at 95 °C for 5 min.

### Potassium permanganate DNA footprinting

DNA was labeled by [γ-^32^P] ATP individually at the 5’ end of the template or the non-template strands. The labeled DNA (~12000 cpm) and 150 nM Eσ^70^ were used to form the paused TECs. The sample was mixed with equal volume of 20 mM KMnO_4_ by vortexting for 15 s and quenched by 1.3 M β-mercaptoethanol. After adding 80 μg salmon sperm DNA (Invitrogen) and nuclease-free water to a total volume of 100 μl, the DNA fragments were extracted by PCI and precipitated by adding 1/10 volume of sodium acetate and 2.5 volumes of ethanol for 1 h at −20 °C. The pellet was resuspended in 10% (v/v) piperidine and treated for 15 min at 90 °C. The DNA fragments were re-precipitated and washed twice with 70% ethanol. The DNA pellet was dissolved in 20 μl nuclease-free water, dried by vacuuming and dissolved in the loading buffer (95% formamide, 20 mM EDTA pH 8.0, 0.2% SDS, 0.05% xylene cyanol and bromphenol blue). The sequencing ladders were generated by a Thermo Sequenase Cycle Sequencing Kit (ThermoFisher). The resultant DNA products were analyzed by 10% (19:1, acrylamide: *bis*-acrylamide) PAGE containing 7.5 M urea.

### RNA-seq and data analysis

To extract total RNA for RNA-seq, 8 ml *E. coli* cells grown to mid-log phase (OD_600_ = 0.5) were spun down, resuspended in 800 μl TRIzol (Invitrogen) and incubated for 4 min at 95 °C. The total RNA was purified by 400 μl PCI extraction and 200 μl chloroform extraction. After centrifugation, an equal volume of isopropanol was added to the top water phase to precipitate the RNA. The genomic DNA was digested with 50 U DNase I for 30 min at room temperature. The RNA was purified by RNeasy Mini Kit (Qiagen) and its concentration was quantified by Agilent 2100 bioanalyzer. The libraries were constructed using TruSeq Stranded Total RNA Library Prep Kit (Illumina) and applied to MiSeq using 2 × 150 bp paired-end sequencing at the Center for Cancer Research Sequencing Facility. The reads that passed quality control and filtering of the raw data were aligned to the *E. coli* genome NC_000913.2 using STAR^55^. The raw counts of the aligned reads for each gene were calculated by HTseq^56^. Fold changes of genes transcription between different samples were calculated by DESeq2^57^.

## Data availability

All RNET-seq and RNA-seq data from this study were deposited to NCBI’s Gene Expression Omnibus (GEO) database under the accession number GSE147611.

## Code availability

The custom scripts used for the analysis of RNET-seq data are available at https://github.com/Mikhail-NCI-Lab/RNET-seq_code.

## Acknowledgements

We thank D. Jin for *E. coli* RNAP and σ^70^, L. Lubkowska for Gre proteins and S. Yang for pCas and pTarget plasmids. We are grateful to T. D. Schneider for helpful discussion and critical reading the manuscript. We also thank the NIH Intramural Sequencing Center and the CCR Sequencing Facility for Illumina sequencing. This work was supported by the Intramural Research Program of the National Institutes of Health, National Cancer Institute, Center for Cancer Research to M. Kashlev.

## Author contributions

Z.S. and M.K. conceived and designed the project. A.Y. optimized the RNET-seq workflow. Z.S. performed the RNET-seq, RNA-seq and biochemical experiments. Z.S., C.M. and P.F. wrote and executed the custom scripts. Z.S. and M.K. analyzed the data. Z.S. and M.K. wrote the manuscript.

## Additional information

Extended data is available for this paper.

Supplementary information is available for this paper.

